# Zero-shot imputations across species are enabled through joint modeling of human and mouse epigenomics

**DOI:** 10.1101/801183

**Authors:** Jacob Schreiber, Deepthi Hegde, William Noble

**Affiliations:** University of Washington, Seattle, Washington, United States

**Keywords:** deep learning, functional genomics, tensor factorization

## Abstract

Recent large-scale efforts to characterize functional activity in human have produced thousands of genome-wide experiments that quantify various forms of biochemistry, such as histone modifications, protein binding, transcription, and chromatin accessibility. Although these experiments represent a small fraction of the possible experiments that could be performed, they also make human more comprehensively characterized than any other species. We propose an extension to the imputation approach Avocado that enables the model to leverage genome alignments and the large number of human genomics data sets when making imputations in other species. We found that not only does this extension result in improved imputation of mouse functional experiments, but that the extended model is able to make accurate imputations for protein binding assays that have been performed in human but not in mouse. This ability to make “zero-shot” imputations greatly increases the utility of such imputation approaches and enables comprehensive imputations to be made for species even when experimental data are sparse.

**CCS CONCEPTS:** • Computing methodologies → Neural networks; Factorization methods; • Applied computing → Bioinformatics; Genomics.

**ACM Reference Format:** Jacob Schreiber, Deepthi Hegde, and William Noble. 2020. Zero-shot imputations across species are enabled through joint modeling of human and mouse epigenomics. In *ACM-BCB 2020: 11th ACM Conference on Bioinformatics, Computational Biology, and Health Informatics, Sept 21–24, 2020, Virtual*. ACM, New York, NY, USA, 9 pages. https://doi.org/10.1145/1122445.1122456

## 1 INTRODUCTION

Genome-wide sequencing-based measurements, such as ChIP-seq for measuring histone modifications and protein binding, RNA-seq for measuring transcription, and DNase/ATAC-seq for measuring chromatin accessibility, quantitatively characterize the molecular basis for cellular mechanisms, differentiation, and disease. Consequently, individual investigators perform these assays to answer specific research questions, and large consortia—such as the Roadmap Epigenomics Consortium [14], the ENCODE Project [5], and the International Human Epigenomics Consortium [2]—perform and collect thousands of them into compendia that broadly characterize functional activity across a variety of primary cells and tissues (“biosamples”).

Unfortunately, these compendia are rarely complete, and this incompleteness is worse in species other than human. As of June 12, 2020, the ENCODE Project portal (https://www.encodeproject.com) hosts only 1,814 epigenomic experiments mapped to the mouse reference genome mm10, in contrast to the 9,135 experiments mapped to the human reference genome hg38. The experiments performed in mouse span fewer assays and biosamples than the human experiments, and each mouse biosample is generally less well assayed than a typical human biosample. Perhaps most importantly, the overall characterization of protein binding is far sparser in mouse than in human, despite proteins, such as transcription factors, playing crucial regulatory roles in the cell. To illustrate this difference in sparsity: the best characterized human biosample, K562, has 558 protein binding experiments mapped to hg38, whereas the best characterized biosample in mouse, MEL, has only 49 protein binding experiments mapped to mm10. Further, only 32 mouse biosamples have been assayed for protein binding at all, whereas hundreds of human biosamples have been assayed for the binding of at least one protein.

To mitigate this incompleteness, several computational methods have been proposed for imputing the signal of experiments that have not yet been performed [4, 7]. A recent method, Avocado [22], is a deep tensor factorization model that treats a compendium as an incomplete 3D tensor with axes corresponding to biosample, assay type, and genomic position. Avocado learns latent representations for each of the three axes independently and, by combining these representations using a neural network, is able to impute the signal for any genomics experiment contained within the tensor. However, current imputation approaches, including Avocado, are restricted to operate on the set of assays and biosamples where at least one experiment has been performed. Because data is available for far fewer assays in mouse than in human, existing imputation methods are limited in two ways: first, there is less available data for training, and second, there are far fewer assays for which even a single experiment has been performed.

We address both of these problems with an extension to Avocado that jointly models mouse and human genomics experiments. In this extension, we expand the data tensor to include assays and biosamples from both species, and use an alignment between the human and mouse genomes (which we refer to as “synteny information”) to map signal from human experiments to positions within the tensor. A side effect of this procedure is that most values within the new tensor will come from human experiments; thus, we modify the optimization strategy for Avocado to sample values from each species with equal probability to prevent the model from focusing too much on human signal. The inclusion of human experiments dramatically increases the amount of data available for training mouse imputation models and, importantly, enables imputations to be made for assays that have only been performed in human. In machine learning terminology, imputing these assays in mouse is an example of a “zero-shot” problem, where a model makes predictions in a setting where it was not given any training data. To our knowledge, no other large-scale imputation approach is capable of making imputations in this zero-shot setting.

A notable aspect of this extension is that it takes advantage of the evolutionary relationship between human and mouse. Mouse and human genomes contain large amounts of shared sequence [3, 24, 27], with approximately 40% of the human genome aligning to the mouse genome [18]. This similarity is the basis for methods that have identified previously unknown regulatory elements by transfering functional annotations across species [10], and also extends to the biochemistry of the cell. Naturally, transcription and accessibility measure the same underlying phenomena in both species, but even histone modifications and proteins are known to play similar regulatory roles. For instance, in both species the histone modification H3K4me3 is enriched in active promoters [1, 9], H3K27me3 is enriched in repressed promoters [19], and the transcription factor MYC is associated with cell growth [16, 17].

We find that our proposed joint optimization procedure allows Avocado to make higher quality imputations for mouse than it does using mouse data alone. When we compare a model trained using our procedure to a model trained using only mouse data, we observe an overall decrease in the mean-squared-error (MSE) of 4.4% and, when partitioning the experiments by activity type, a decrease in MSE of 24% for transcription-measuring experiments. Further, we show that this improvement requires both human experiments and synteny information, and that simply incorporating human experiments is not as effective. Interestingly, we found that some assays exhibited improved performance in mouse even when no experiments of that type had been performed in humans. In these cases, the improvement likely arose because Avocado identified similarities between assays using only the mouse experiments and then using the additional human experiments for those assays to boost performance.

We next demonstrate that this procedure allows for imputation of assays that have been performed in human but not in mouse. Even though these experiments could not be imputed using traditional methods, we show that the resulting imputations are highly accurate, with almost one-third of proteins showing a 50% decrease in MSE compared to the strong average activity baseline and two-third showing at least 20% decrease in MSE. Although we did not find a strong correlation between the amount of available human data for an assay and improvement in performance, we did find that the our approach exhibited the strongest improvements over the baseline in the most difficult cases. We find that that a major source of improvement in these zero-shot imputations is that, despite predicting a similar number of peaks as the baseline approach, the peaks occur more frequently within accessible chromatin.

## 2 METHODS

### 2.1 Synteny mapping

A critical step in our procedure is mapping signal from human experiments to positions on the mouse genome. We build this mapping using sequence alignment annotations from the hg38ToMm10 liftOver file available at http://hgdownload.cse.ucsc.edu/goldenpath/hg38/liftOver/. The liftOver file consists of “chains” that define gapped pairwise alignments between the two species. In the file, each chain is described by a single header line followed by one line for each ungapped sequence alignment. When the alignment for one of the species occurred on the minus strand, the positions were adjusted appropriately, i.e. counting is done with respect to the end of the chromosome instead of the beginning. When building our mapping, we used only these ungapped alignment, rather than the entirety of the chain, to diminish the effect of gaps. It is worth noting that the hg38ToMm10 and mm10ToHg38 liftOver files are not symmetric because the liftOver tool contains only unique coverage for the target species but not the query species. We intentionally chose the hg38ToMm10 file because we wanted to include as many connections from the mouse genome to the human genome as possible.

Next, we convert these sequence alignments to alignments of 25 bp bins because that is the resolution Avocado operates at. First, ungapped alignments that are shorter than 25 bp are discarded because they do not span a single bin and would distort the signal in cases where the alignment is split by the boundaries of a bin. For the remaining ungapped alignments, the starts and ends are divided by 25 and rounded down (integer division) for each species independently to get an alignment of bins. For example, an ungapped alignment between positions 120–215 of mm10 and 72–167 of hg38 is converted to an alignment of zero-based half-open intervals of bins 4–8 for mouse and 2–6 for human. The output of this step is a triplet of arrays for each mouse chromosome: positions on the mouse chromosome with aligned sequence, chromosomes in the human genome that the mouse position aligned to, and positions on the respective chromosome in the human genome that aligns to the mouse position.

### 2.2 Data sets

In total, we downloaded and processed 8,015 genomics experiments hosted on the the ENCODE project portal (https://www.encodeproject.org). These experiments included 6,870 measuring biochemical activity in humans, denoted as ENCODE2018-Full, and 1,145 measuring biochemical activity in mouse, denoted MouseENCODE2019.

The output from these experiments was processed in a similar fashion to previous work involving Avocado [21, 22]. The sequencing reads were processed using the ENCODE Processing Pipeline [15] and mapped to either human genome assembly hg38 or mouse genome assembly mm10. The resulting signal values are − log 10 p-values for the ChIP-seq and ATAC-seq data, read-depth normalized signal for DNase-seq, and normalized stranded read coverage for the RNA-seq experiments. When a pooled replicate was present for an experiment we preferentially chose it; otherwise, we chose the second replicate if two replicates, but not a pooled version, were present, and the first (and only) replicate otherwise.

After these experiments, the data were then further processed before model training. First, the signal was downsampled to 25 bp resolution by taking the average signal in each non-overlapping 25 bp window. Second, an inverse hyperbolic sine transformation was applied to the downsampled data. This transformation has been used previously to reduce the effect of outliers in epigenomic signal [4, 11]. The inverse hyperbolic sine function is similar to a log function except that it is defined at 0 and is almost linear in the range between 0 and 1. The transformed tracks are used both for training and evaluating the models.

Finally, we used our synteny mapping to extract measurements from human experiments that mapped to each mouse chromosome. This procedure was fairly straightforward and involved simply applying the triplet of arrays to, using the first array, define the position on the mouse genome and, using the second and third arrays, copy the signal from the corresponding positions on the human genome. When representing the data as a tensor, the measurements from human experiments appear as partially complete blocks attached to the tensor of mouse measurements at positions where the mouse genome aligns to a region on the mouse genome (Figure 2A).

**Figure 1:**
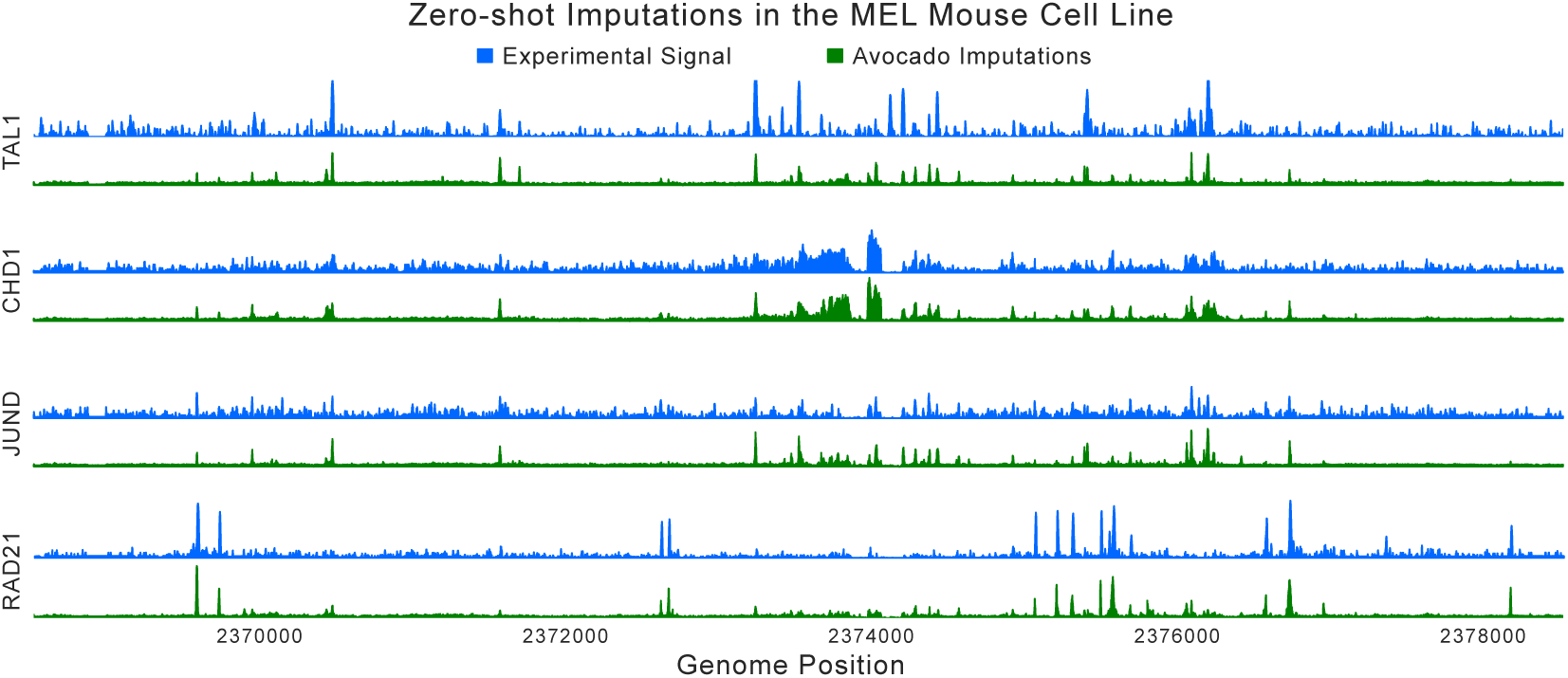
The experimental ChIP-seq signal (in blue) and corresponding zero-shot imputations (in green) made at 25bp resolution for four proteins in the mouse cell line MEL using a model trained with the procedure we propose in this work. All tracks depict signal at the same genomic locus and are on the same scale.

**Figure 2:**
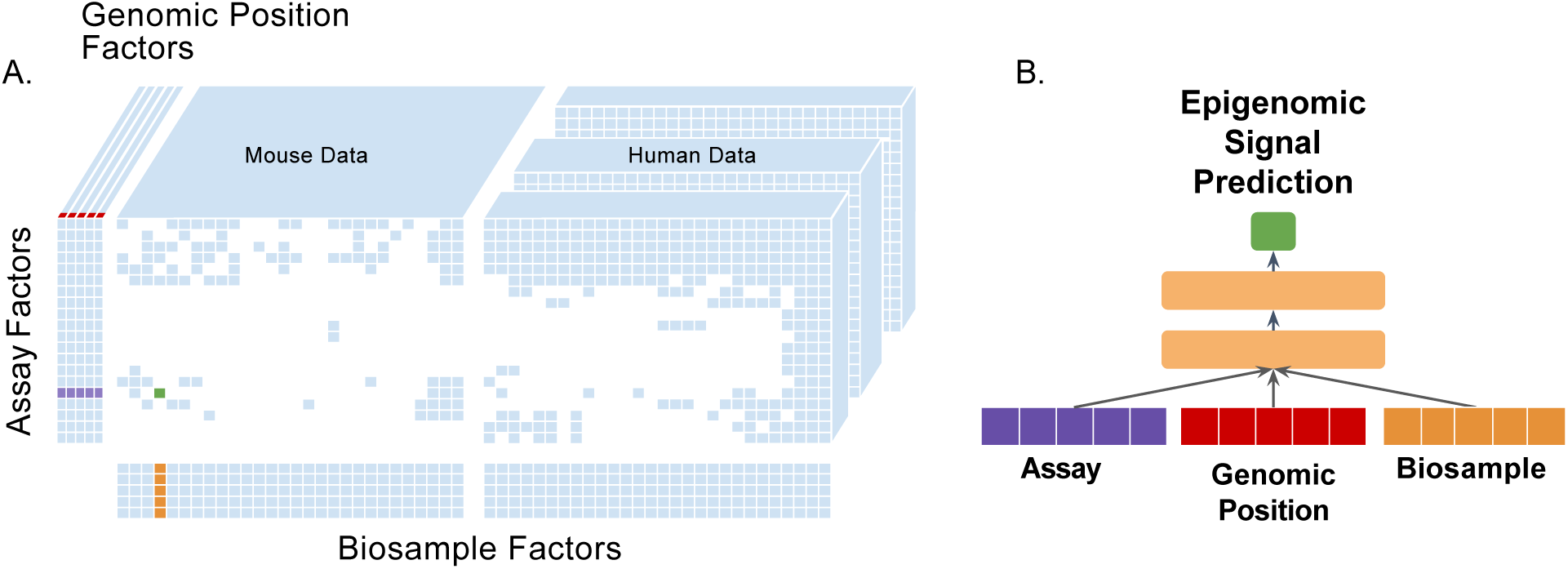
The cross-species Avocado model. (A) A schematic of the expanded data tensor with shared assay and genomic position axes. Mouse data is available for all positions on the mouse genome, whereas human data is only available for positions in the mouse genome with aligned human positions. (B) The neural network takes in factor values for a single biosample, assay, and genomic position, and predicts the corresponding value within the tensor.

### 2.3 Avocado model

Avocado is a deep tensor factorization method for modeling a compendium of experiments as a partially-filled 3D tensor [21, 22]. The model is comprised of latent representations for each of the three dimensions of the tensor—biosamples, assay types, and genomic positions— (Figure 2A) and a neural network that combines these representations in a non-linear manner to predict values within the tensor (Figure 2B). The assay representations have 256 dimensions, the biosample representations have 32 dimensions, and the genomic representation is split between three resolutions with 25 dimensions at 25 bp resolution, 40 dimensions at 250 bp resolution, and 45 dimensions at 5 kbp resolution. The neural network takes the concatenation of these representations as input, has two dense layers of size 2,048 with ReLU activation functions, and outputs the signal at one position for a single experiment. The latent representations and the weights of the neural network model are trained jointly using the Adam optimizer [13] with default hyperparameters. Avocado can impute any genomics assay within the tensor by sequentially substituting in the entire set of genome factors.

Although the topology of Avocado remains the same in our extension, we made two changes to accomodate the expanded data tensor. The first change is that the assay axis contains the union of all assays in both species, and the biosample axis contains the union of all biosamples in both species. During training, an example drawn from a mouse experiment involves a triplet as input specifying the genome position, the assay, and the mouse biosample. Similarly, an example drawn from a human experiment involves a triplet specifying the aligned position on the mouse genome, the assay, and the human biosample. The second change involves modifying how data are sampled during training. In our extension, half of the values in each batch comes from each species. We made a third change that was not critical for our procedure but empirically improved the convergence of our model: instead of training on genomic positions sequentially, as in previous work, we permute the order that genomic positions are sampled each epoch.

As an additional baseline, we construct an Avocado model that incorporates human experiments but does not use synteny information. This model involves constructing a tensor of measurements where each of the three axes is the concatenation of the mouse and human elements. The primary difference between this and our proposed extension is that this model will have a separate set of factors for the mouse and human positions, instead of having only genomic factors for mouse positions. This model is trained using the same data as the model that incorporates synteny information, i.e. measurements from the portions of the human genome that align to the mouse genome, but this alignment information is discarded. Thus, in the data tensor, there is only an overlap in assays between experiments from the two species.

### 2.4 Calculation of average activity

We use the term “average activity” to refer to the average signal value for each assay across a set of training tracks at each position in the genome. This value can be used as a strawman imputation procedure. As a baseline, the average activity score is typically much stronger than the simple average signal value across all loci, which is a more traditional baseline. Generally, we calculate the average activity for an assay by calculating the average signal across the experiments in the training set used to train the model we are evaluating. In the context of three-fold cross-validation, for each fold the average activity is calculated for each assay using the experiments in the other two folds (the training set). Formally, the average activity *AA* for an assay *a* from the set of all training set experiments of t t assay *E* at position *i* in the genome is calculated as 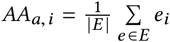

### 2.5 Model evaluation

We evaluate the models presented in this work using the mean-squared-error across all folds in a cross-validation that partitions entire experiments into three folds. Due to computational limitations that arise when training many models for comparison, we limit our evaluations to chr3, chr11, and chr19 of the mouse genome. We chose chr19 because it was the smallest, and the other two were chosen randomly. Because the training process involves fitting each chromosome independently, the released model is available for the entire mouse genome.

For our initial cross-validation experiment in Section 3.1, we partition experiments into folds that attempted to include each assay at least once in each fold. Some assays had been performed fewer than three times, and so experiments for these assays were partitioned into folds normally for training but were excluded from evaluation. This reduced the number of mouse experiments from the 1,145 experiments used for training to 1,116 used for evaluation.

Our evaluation of the zero-shot imputations involved a cross-validation of protein binding assays instead of experiments. Each step of cross-validation involved filtering out all experiments for the assays in the specified fold, training a model using all remaining human and mouse data, and evaluating performance solely on those experiments that had been filtered out. While this validation did not require that an assay had been performed multiple times, like the initial cross-validation did, the same set of 1,116 were used for evaluation because the average activity baseline that we compared against required multiple experiments to be performed per assay.

## 3 RESULTS

### 3.1 Joint optimization improves imputations in mouse

Our primary hypothesis is that an imputation approach that jointly models experiments from multiple species will perform better, and be more comprehensive, than an approach that models experiments from only a single species. Accordingly, we began by quantifying the benefit that including human experiments in the training process had when imputing mouse experiments. Our first evaluation was a three-fold cross-validation, where the three folds came from splitting the experiments in MouseENCODE2019 into three partitions such that each assay had been performed at least once in each partition (see Methods for details).

We evaluated three Avocado models that were trained using differing amounts of information. The first model was trained using only mouse experiments and so represented model performance on the standard imputation task. The second model was trained using both mouse and human experiments, but without synteny information, by simply concatenating together the two tensors along the genome axis (see Methods for details). The third model is trained on the same set of experiments as the second model but uses our joint optimization procedure that accounts for synteny. We train these three models so that we can separately assess the benefit of including human genomics experiments and the synteny information. Importantly, when performing cross-validation, models that include human experiments are given access to the entirety of the human data sets in each fold. Although the second and third models were trained using both human and mouse epigenomic experiments, all three models were only evaluated on their ability to impute mouse epigenomic experiments.

As a baseline for the imputation approaches described above, we calculated the average activity of each assay (see Methods for details). The average activity for an assay is the average signal at each position exhibited by the training set experiments that are of that assay. The average activity baseline represents a simple rule that regions of consistently high or low signal in the training set will exhibit similar behavior in the test set [23]. Accordingly, improvement over the average activity baseline generally indicates the prediction of cell type-specific activity.

We comprehensively calculated the performance, as measured by MSE, of each of the models using cross-validation. Overall, we found that the more information the model had access to, the better the model performed: the model that used human data but not synteny information outperformed the mouse-only model (paired t-test p-value of 9.39e-8) but itself was outperformed by the model that used both (p-value of 6.54e-7, Table 1). The largest performance improvement came from imputing histone modification experiments (p-value of 1.86e-28), which were plentiful in both human and mouse contexts. Proportionally, the largest improvement is in the experiments that measure transcription (p-value of 8.24e-6), likely because the human data sets contain a large number of transcription-measuring experiments. A similar trend—a large improvement in predicting transcription in mouse when leveraging human data sets—has also been observed by Kelley [12] when predicting functional state using nucleotide sequence alone.

**Table 1:** Imputation performance. The MSE computed both overall across all experiments and for each of the four main forms of biological activity in MouseENCODE2019. For each measure, the score for the best-performing model is in boldface if the improvement over the Mouse Only model is significant (paired t-test p-value ≤ 0.01). For transcription experiments, the two models that use human data sets are not significantly different from each other (p-value of 0.363) and so both numbers are bolded.

| MSE | Overall | Histone Modifications | Protein Binding | Transcription | Accessibility |
| --- | --- | --- | --- | --- | --- |
| Average Activity | 0.0824 | 0.1056 | 0.0942 | 0.00217 | 0.0519 |
| Mouse Only | 0.0596 | 0.0763 | 0.0646 | 0.00194 | 0.0440 |
| + Human Data Sets | 0.0582 | 0.0741 | 0.0654 | **0.00151** | 0.0433 |
| + Synteny Information | **0.0569** | **0.0721** | 0.0653 | **0.00148** | 0.0435 |

We identified two sets of loci with distinct characteristics where our approach led to improved performance. In the first set of loci, each imputation method generally outperformed the average activity baseline, but our approach led to more accurate predictions of the exact signal values than the mouse-only model (Figure 3A). The second set of loci were characterized by similarly poor performance of both the average activity baseline and the mouse-only model (Figure 3B), but better performance using our proposed approach. These loci were prominent among transcription-measuring experiments, and might explain the large proportional gains in performance observed when using our method.

**Figure 3:**
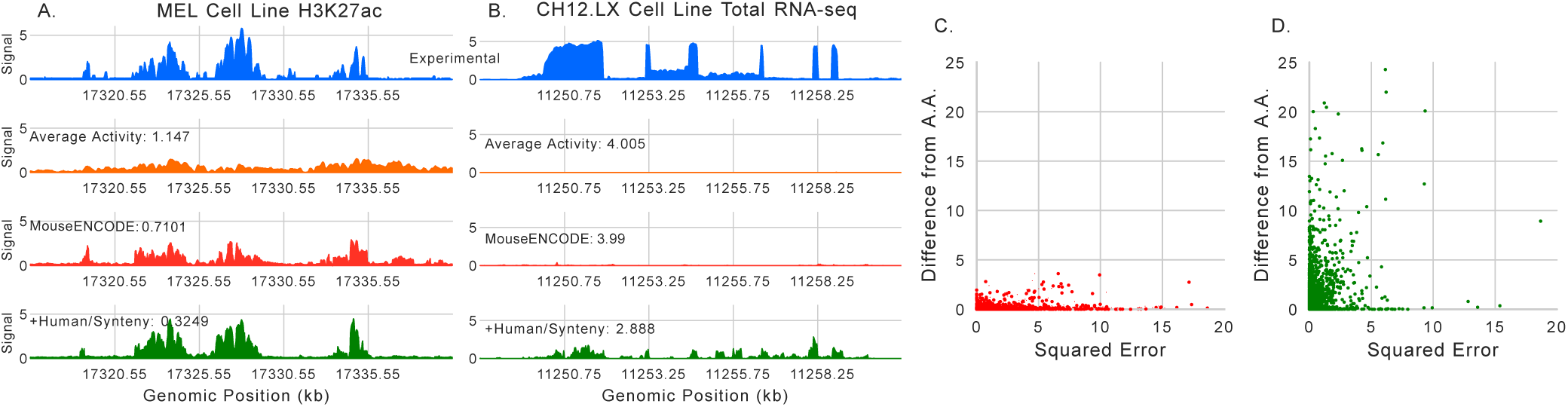
Examples of real and imputed signal. (A) An example of experimental signal for H3K27ac in the MEL cell line (blue), the average activity (orange), the imputed signal from a model trained using only data from mouse (red), and the imputed signal from a model trained using our procedure (green). Each approach is annotated with the MSE compared to the experimental signal for the visualized region. (B) The same as (A) except for total RNA-seq in the CH12.LX cell line. (C) The MSE between the imputations made from the mouse-only model and the experimental signal versus the difference between the imputed signal and the average activity, from all positions exhibing non-zero experimental signal in chr3 for the experiment in (B). (D) The same as (C) except for the model trained using our procedure.

To investigate the second set of loci further, we analyzed all positions with non-zero experimental signal in chr3 for a total RNA-seq experiment performed in CH12.LX. We found that the MSE for these positions decreased from 0.191 when only the mouse-only model to 0.09 when using our approach, but that the MSE between the imputations and the average activity increased from 0.017 when using the mouse-only model to 0.20 when using our approach (Figure 3C/D). These results indicate that a source of error for the mouse-only model is making imputations that too closely resemble the average activity, i.e. are not biosample specific, and that the additional information provided to our approach can improve the biosample-specificity of the resulting imputations.

Interestingly, we found that, for two of the three assays with the largest absolute gain in performance (MYOG and H3ac, with decreases of 8% and 12% MSE respectively), no experiments had been performed in human. This observation may initially appear counterintuitive. However, we note that both MYOG and H3ac co-occur with other biological activity that has been measured in both mouse and human. Specifically, according to STRING-DB [25], MYOG interacts with five proteins that have been measured in both species, most prominently TCF12 with five experiments in human, and H3ac generally coincides with H3K4me3. The improved performance on MYOG and H3ac indicates that our extension can benefit the imputations not only of assays that have been performed in human, but also those whose activity is correlated with assays that have been performed in humans.

### 3.2 Joint optimization enables zero-shot imputations

Encouraged that our proposed procedure led to an improvement in overall performance, we hypothesized that the same procedure could allow Avocado to make predictions in mouse for assays that have only been performed in human. We refer to this as the “zero-shot” setting because the model has no training data in mouse for the assays that it is making imputations of. Although there are a variety of assays where data is available in human but not in mouse, we focused our evaluations here on imputing protein binding experiments because of the importance that protein binding plays in gene regulation and because many proteins have had their binding characterized in human but not in mouse.

We began by investigating model performance in two related zero-shot settings. In the first setting, protein binding assays (not the experiments themselves) were divided into three partitions, and cross-validation folds were constructed such that all experiments in mouse from assays included in the corresponding partition were removed. This resulted in three folds of experiments where each fold excluded all mouse experiments from one-third of protein binding assays. In the second setting, all protein binding experiments performed in mouse were excluded. Although the first setting is more realistic, because some protein binding experiments have already been performed in mouse, the second setting allows us to investigate the performance benefit of including these experiments when making zero-shot imputations. We compared performance in both of these settings to the same average activity baseline from the previous section. This baseline is even more difficult to beat in these zero-shot settings because it is explicitly derived from data that the model does not have access to, i.e., protein binding experiments in mouse.

Despite the strength of the average activity baseline in this context, we found that the Avocado models trained using our procedure generally outperformed it. Visually, we observed that the imputations in both settings were similar to the experimental signal (Figure 4A/B). Overall, in the first setting, the MSE dropped significantly from 0.919 when using the average activity to 0.778 when using our procedure (paired t-test p-value of 3.29e-5) and, in the second setting, remained at 0.919 (p-value of 0.996, Figure 4C). In both settings, models that leveraged the synteny information significantly outperformed those that did not (p-values of 5.07e-5 and 7.68e-6 respectively). The improvements in performance when using synteny are over 1 and 2 orders of magnitude larger (respectively) in the two zero-shot settings (0.0045 and 0.0246) than in the original cross-validation experiments (0.00017), indicating that synteny information is particularly useful for identifying protein binding sites in the absense of experimental data sets.

**Figure 4:**
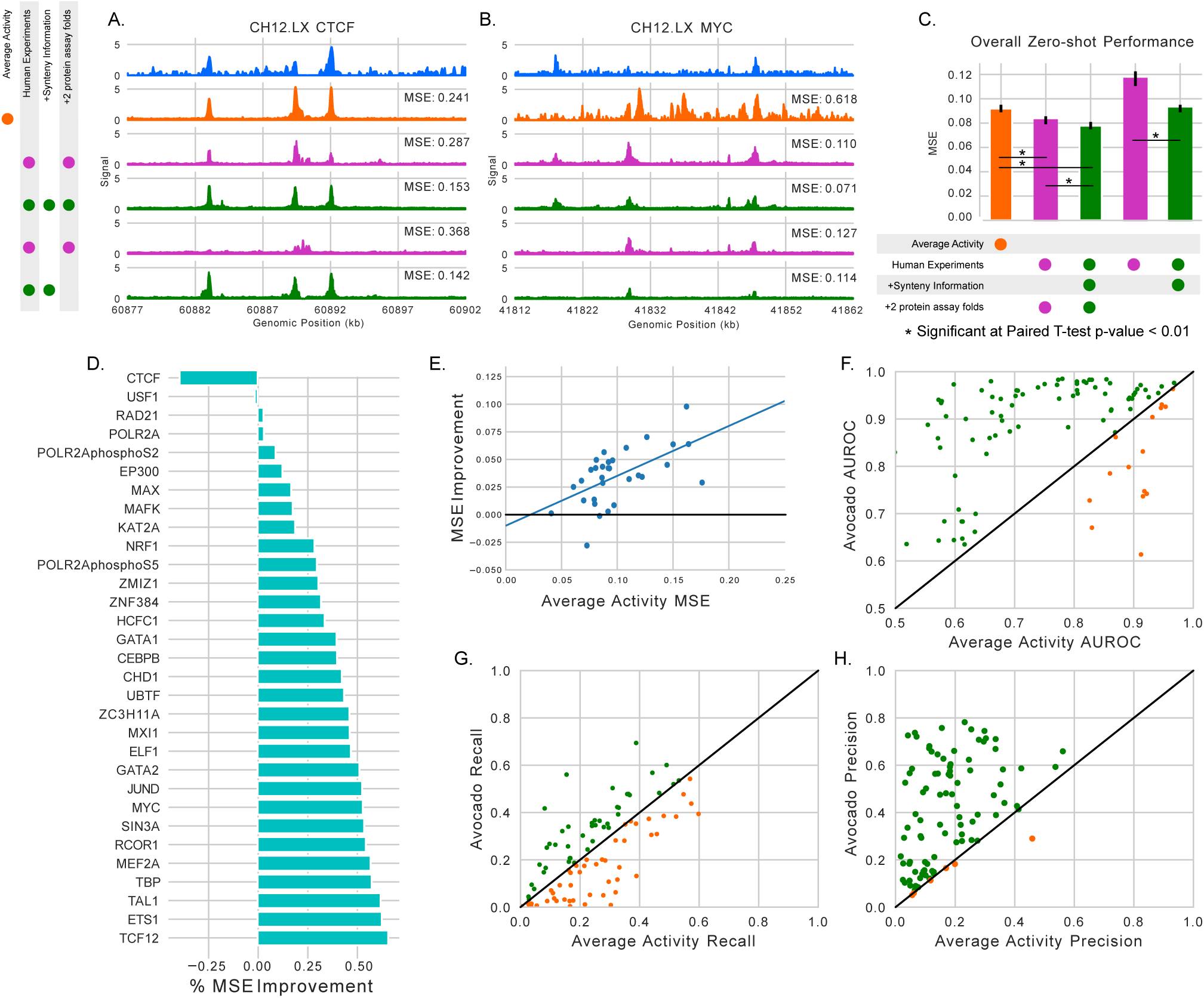
Examples and evaluation of zero-shot imputations. (A) The experimental signal, average activity, and imputed signal of four models for binding of CTCF in CH12.LX. The MSE for each approach compared to the experimental signal in the displayed window is also shown to the right. The legend to the left shows the set of experiments used to train each model. (B) The same as (A) but for the binding of MYC in CH12.LX. (C) The overall performance of each approach with four statistically significant relationships highlighted. (D) The percentage improvement between imputations from our proposed methodology (that uses two folds of protein binding experiments) and the average activity. (E) The relationship between the performance of the average activity baseline and the absolute improvement that our approach exhibits with a best-fit line drawn. (F) The AUROC from identifying DNase peaks using imputed protein binding signal and the average activity. (G) The recall for the same task as (F) when the protein binding signal is binarized with the same threshold as the DNase signal. (H) The same as (G) except reporting precision instead of recall.

Next, we evaluated the performance of the average activity and our zero-shot imputations on a per-protein basis. We found that our imputations improved upon the average activity baseline for 29 out of 31 proteins (Figure 4D). This improvement was often substantial: for example, our models decreased the MSE by at least 50% relative to the average activity for almost one-third (10) of proteins and decreased the MSE by at least 20% relative to the average activity for over two-thirds (22) of proteins. The average decrease in MSE was 0.035 and, on average, each protein saw a 33.7% improvement relative to the average activity baseline. These results are particularly striking because the model had not been explicitly exposed to nucleotide sequence, information about motif occurrances, or examples of the respective protein binding in other mouse biosamples.

However, we noted that the model does not improve over the average activity baseline for all proteins. Specifically, our model underperforms the average activity baseline on CTCF and USF1 (though we note that the performance difference on USF1 is very small). It is not entirely surprising that the average activity baseline is strong for CTCF. First, CTCF has been performed 33 times in mouse, almost twice as many times as the next leading assay POLR2A at 19 and over 8 times more than the average (4.1 times), and so the average activity is likely a more robust estimator. Second, CTCF binding is generally very similar across biosamples. This means that a baseline derived from observed binding in some biosamples will be a very strong baseline for prediction in a new biosample, particularly when compared to a model that does not observe this binding even a single time.

We then sought to better understand the cases where our model achieved large gains. A reasonable hypothesis is that the performance of our approach, which relies on human data to make zero-shot imputations, is associated with the amount of available human data. However, we did not find strong evidence that this was the case. In fact, we initially found a negative correlation between the number experiments performed in human and the resulting improvement in performance in mouse (*r* = −0.57). This negative correlation was primarily driven by four outlier proteins, including CTCF, that had been assayed in humans significantly more than the norm (between 45 and 194 times compared to 11 times or fewer for all other proteins). When we excluded these four we found that the correlation decreased (to *r* = −0.11). In contrast, we found that the performance of the average activity baseline itself was much more correlated with performance improvement, with a correlation of 0.579 over all proteins and 0.697 when excluding the four outliers (Figure 4E). This finding is not unexpected: the worse the average activity baseline, the larger the room for improvement. However, confirming that our method exhibits larger improvements in more difficult cases is still worthwhile.

Finally, we evaluated the biological consistency of these imputations by measuring how well protein binding coincided with chromatin accessibility. Accordingly, we identified 95 protein binding experiments from mouse biosamples that also had a DNase-seq experiment performed in them, and we measured the extent that imputed signal for each protein was localized within experimental DNase-seq peaks. This measurement was, specifically, the area under the receiver-operator-characteristics curve (AUROC) for the classification problem of identifying DNase peaks, defined by having a read-depth normalized signal value above 2, using the protein binding signal. The AUROC has the intuitive interpration of being the probability that signal is higher within accessible sites than elsewhere. Overall, we found that Avocado’s imputations had a higher AUROC than the average activity baseline in 79 of the 95 experiments we considered, and had an average improvement of 0.119 AUROC (Figure 4F).

We reasoned that there were two possible sources for this improvement: imputed peaks could simply be occuring at all accessible sites in each biosample, or more of the imputed peaks could fall within accessible sites. Fortunately, distinguishing between these sources is easy because they correspond simply to recall (proportion of accessible loci that exhibit imputed peaks) and precision (proportion of imputed peaks that occur at accessible loci) respectively. The first source would indicate undesirable behavior from the model, in that the model cannot identify which proteins bind at which accessible loci, whereas the second would indicate desirable behavior, because the model would not predict binding at inaccessible loci.

When we calculated these values we found that the this increase in AUROC was driven largely by increases in precision but not in recall. Specifically, we found that the average activity had a recall that was only larger by 0.004 (p-value of 0.769, Figure 4G), whereas the imputations had a precision that was larger by 0.23 (p-value 1.2e-18, Figure 4H). The similar recall scores indicate that protein binding predictions from both approaches cover roughly the same number of accessible loci, but the higher precision score for the imputations indicates that the imputed signal is more concentrated in accessible loci than the average activity is, i.e. that fewer peaks occur outside accessible loci. Taken together, these results confirm that our protein binding imputations are concentrated at accessible sites but do not simply associate accessibility with protein binding.

## 4 DISCUSSION

In this work, we introduce an extension to the imputation method Avocado that improves its performance by jointly modeling experiments from multiple species. We find that this extension improves the performance of a mouse imputation model, and that part of this improved performance comes from imputing biosample-specific signal at loci where a mouse-only model simply imputes the average activity. Encouragingly, we find that improvements occurred even for assays that have not been performed in human, demonstrating the broad impact that modeling experiments from multiple species can have. We then show that our extension enables the model to make imputations in mouse for assays that have only been performed in humans, and that these imputations are biosample specific and significantly more accurate than the average activity—a strong baseline for this setting. Altogether, our model is capable of making imputations for over 750 different protein binding assays, most of which have not yet been performed in mouse.

We anticipate that these imputations will be useful in a variety of situations. Naturally, any existing application of imputed data, e.g. augmenting genome segmentation methods [6, 7] or prioritizing experimental efforts [20], will immediately benefit from improved accuracy; however, these applications may further benefit because the imputations that we provide span a much larger set of assays than is covered by available experimental data. Potentially, one could apply a method like the one proposed by Wei et al. [26] to the zero-shot imputations to determine an order that assays should be performed for the first time.

The finding that imputation performance for a particularly assay is not strongly associated with the number of times the assay had been performed in human is important. In particular, it suggests that one does not need to perform an assay in several human biosamples before zero-shot imputations can be trusted. However, this finding may be confounded by the presence of other proteins that bind at a similar set of loci (either because they are co-factors, members of the same protein family, or otherwise). It may be the case that accuracy of protein binding imputation is associated with the number of times that co-binding assays have been performed, but we were not aware of a simple way to determine this.

We intentionally do not consider the binding patterns of proteins when constructing folds for our zero-shot imputation setting. As a result, some proteins with similar binding patterns may occur in different folds of cross-validation. We reasoned that this approach for constructing folds is not problematic for evaluation because, realistically, not all proteins with similar binding patterns will be assayed together and, for the task of imputing protein binding, one would expect a method to make use of close relationships. Indeed, one would be disappointed if a model trained using MYC experiments exhibited extremely poor performance at imputing the binding of its near homolog, MAX. That being said, it is not unreasonable to initially be concerned about the generalization capabilities of the model to proteins whose binding patterns are completely different from those already performed in mouse. For those proteins, we expect that the performance of the model trained using no protein binding experiments at all in mouse represents a lower bound of performance.

Although this work focuses on modeling mouse and human experiments, the method we propose is general to any pair of species. It is likely that the performance of the model will be related to the evolutionary distance between the species: when this distance is high, the assumption that our model makes about biochemical similarity in the cell may become a source of error in the imputations [8]. For such pairs of species, it may become necessary to increase the flexibility of the model by allowing it to learn species-specific assay representations that are regularized to be similar to each other. On the other hand, careful analysis of errors made by the current approach may yield a data-driven way of identifying biochemistry that differs between species, and where these differences occur along the genome. Regardless, it will be important to keep these assumptions in mind when applying this extension.

Imputations, models, and latent factors produced by this project will be made freely available at https://github.com/jmschrei/avocado.

## ACKNOWLEDGMENTS

This work was funded by NIH award U01 HG009395.

